# The need for alternative plant species interaction models

**DOI:** 10.1101/2020.11.08.373068

**Authors:** Christian Damgaard, Jacob Weiner

## Abstract

The limitations of classical Lotka-Volterra models for analyzing and interpreting competitive interactions among plant species have become increasingly clear in recent years. Three problems that have been identified are (1) the absence of frequency dependence, which is important for long-term coexistence of species, (2) the need to take unmeasured (often unmeasurable) variables influencing individual performance into account (e.g. spatial variation in soil nutrients or pathogens) and (3) the need to separate measurement error from biological variation. We modify the classical Lotka-Volterra competition models to address these limitations and we fit 8 alternative models to pin-point cover data on *Festuca ovina* and *Agrostis capillaris* over 3 years in a herbaceous plant community in Denmark, applying a Bayesian modelling framework to ascertain whether the model amendments improve the performance of the models and increase their ability to predict community dynamics and therefore to test hypotheses. Inclusion of frequency dependence and measurement error improved model performance greatly but taking possible unmeasured variables into account did not. Our results emphasize the importance of comparing alternative models in quantitative studies of plant community dynamics. Only by comparing alternative models can we identify the forces driving community assembly and change and improve our ability to predict the behavior of plant communities.

## Introduction

Plants are sedentary, and neighboring plants affect each other’s growth. The most important effect of growing together with neighboring plants is competition for resources, e.g. light, water and soil nutrients that are necessary for plant growth (Goldberg et al. 1990). However, other mechanisms of plant-plant interactions, such as facilitation in harsh environments, physical interference of vegetative parts and the modification of the behavior of herbivores, pathogens and pollinators, may also play important roles in the assembly of plant communities. Harper (1977) made a comprehensive list of possible negative effects of neighboring plants. As a group, these different mechanisms of negative plant-plant interactions have been called “competition in the broad sense” (Weiner 1993), corresponding to the general definition of competition in ecology as an interaction that is negative for both species (or individuals).

The classical ecological models for exploring the possible effects of competition at the level of the plant community are Lotka-Volterra-type competition models, where the negative effect of neighbors increases linearly with local density (Barabás et al. 2018; Chesson 1994). These models have provided insights into the theoretical conditions necessary for species coexistence at equilibrium. However, it is widely appreciated that frequency-dependent species interaction, in which rare plant species are favored over more common species in their reproduction, growth and/or mortality, may play an important role in plant species co-existence and community dynamics in many plant communities (Chisholm and Fung 2020; Connell et al. 1984). The Janzen-Connell pattern of seedling survival, where seedling survival is reduced close to the parent tree due to herbivores or pathogens, was originally observed in tropical tree species (Wright 2002). However, species-specific soil-plant interactions have received increasing interest as a potentially important and general mechanism for regulating plant populations by hindering local establishment and growth of conspecific plant species in the next generation (Heinen et al. 2020; Mazzoleni et al. 2015a; Mazzoleni et al. 2015b; van der Putten et al. 2013).

In empirical studies of plant communities, the specific mechanisms of plant-plant interactions that influence plant growth and their importance are not known and may vary with the season and size of the competing plants (Gurevitch et al. 2006). Furthermore, it is often not feasible to collect ecological and plant physiological data with sufficient detail to determine the importance of the different mechanisms, let alone model the underlying mechanisms that regulate plant growth.

Most empirical modelling studies of plant species interaction fit Lotka-Volterra-type competition models (e.g Adler et al. 2018; Damgaard 1998; 2005; Law and Dieckmann 2000). This choice is in part motivated by the simplicity of a linear effect of density, ignoring higher-order interactions (Barabás et al. 2018; Chesson 1994). There has been increasing awareness of the limits of Lotka-Volterra-type competition models for understanding inter-specific plant interactions (Mayfield and Stouffer 2017; Neill et al. 2009). For example, if frequency-dependent effects are common and important, they need be included in models of plant community dynamics (Berendse 1979). Alternative approaches are needed and are appearing. For example, Neill et al’.s (2009) model of consumer-resource dynamics based on simple maximum entropy assumptions showed dynamics resembling frequency-dependent species interaction models.

It may be useful to model observed plant population dynamic patterns with several different species interaction models to determine which type of interaction function fits the data best. This would allow us to estimate the relative importance of different types of neighbor interaction mechanisms and to make better ecological predictions of plant community dynamics. The testing of alternative models, rather than one model vs. a null model, is the key to stronger scientific (Platt 1964) and statistical (Gelman et al. 2014) inferences.

For example, consider an interspecific Lotka-Volterra type competition model of population growth in a plant community,

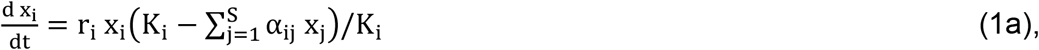

where x is a measure of abundance, r is intrinsic growth rate in the absence of competition, α_ij_ the competitive/facilitative effect of species j on species i (α_ii_ = 1) and K is the carrying capacity. However, if we want to test for the possible effect of frequency-dependency, then this may be modelled by combining the effect of a linear Lotka-Volterra type competition model (eqn. 1a) with a term that models the additional effect of frequency-dependency. Such combined plant-plant interaction effects can be modelled as

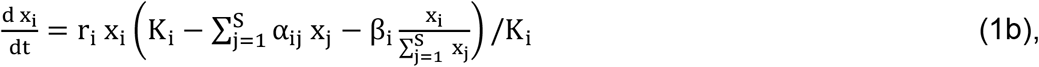

where β > 0 models the negative effect of being common.

The above-mentioned uncertainty about the underlying ecological mechanisms that govern plant inter-specific interactions and which species interaction model is best suited to model empirical competitive growth data is further complicated by the possible role of *unmeasured variables* that influence individual performance and *measurement errors*.

In a recent study, Rinella et al. (2020) demonstrated the possible role of unmeasured variables that are important for plant competitive growth (e.g. small-scale spatial variation in soil nutrient levels or pathogen pressure) in the analysis of inter-specific interactions. Generally, if an unmeasured variable either has an overall positive (e.g. more nutrients) or negative effect (e.g. more pathogens) on plant performance at both the early and later stages of plant development (Fig 1a), then the estimated competitive effect from neighboring plants will be biased (Rinella et al. 2020). This result has received some attention in the statistical literature, where such unmeasured variables are known as instrumental variables or confounding factors (Gelman et al. 2014), and has also been discussed in the plant ecological literature (e.g. Freckleton and Watkinson 2001). The effect of unmeasured variables has rarely been taken into account when fitting data on plant competition in communities.

**Fig. 1.**
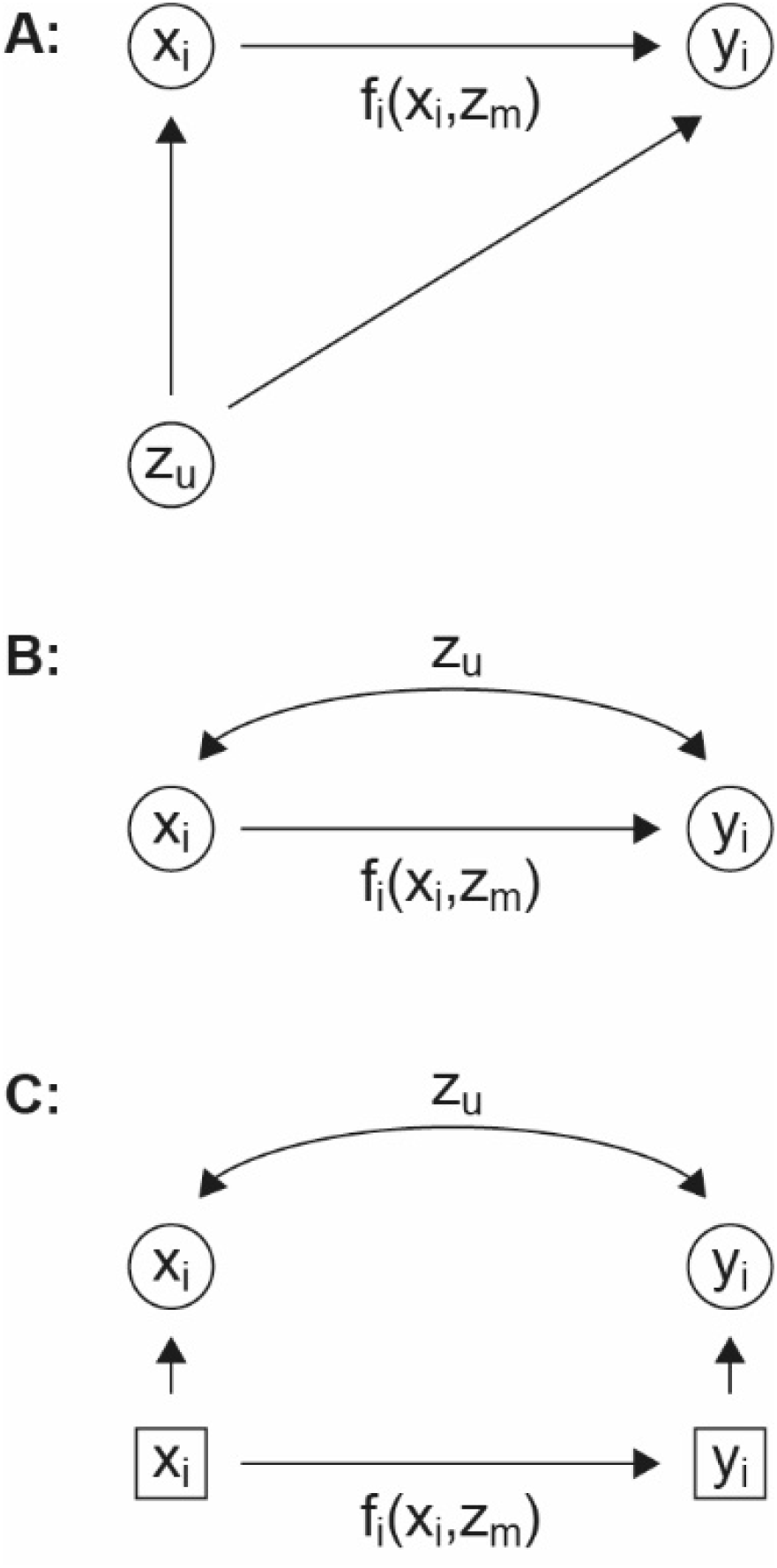
The effect of an unmeasured variable (*z*_*u*_) on individual plant performance at both an early (*x*_*i*_) and later growth stage (*y*_*i*_) of plant species *i*. The competitive growth process is modelled by the function, *f*_*i*_, which depends on *x*_*i*_ and possibly on some measured environmental variables (*z*_*m*_). B: The effect of the unmeasured variable is modelled as the part of the covariance between the early and later growth stages that is not explained by the independent factors in the competition model. C: Hierarchical model in which true but unknown factors affecting plant performance are modelled by latent variables (squares) and measurement errors are separated from process errors. Data are denoted by circles.

The best way to avoid potential confounding effects of unmeasured variables, is to replicate the competition experiments in a relatively homogenous environment. This is not always possible or even desirable (e.g. Freckleton and Watkinson 2001). Rinella et al. (2020) suggested modelling the possible covariance between the early and later growth stages in order to estimate the potential importance of confounding factors that influence individual growth. Since the potential instrumental variables often are difficult or impossible to measure, and it is impossible *a priori* to ensure that all the relevant variables have been measured, then a practical alternative is to model the part of the covariance between the early and later growth stages that is not accounted for by the independent factors in the competition model (Fig 1b). This covariation will at least partly be due to unmeasured variables that affect both the early and later growth stages in the same direction. By including this covariance in the model, the estimated competitive effect of neighboring plants will be less biased.

It has also been demonstrated that sampling and measurement errors may lead to important model and prediction bias, a phenomenon known as regression dilution (Carroll et al. 2006; Damgaard 2020; Detto et al. 2019). We address the possible effect of regression dilution on conclusions inferred from observed inter-specific interactions by fitting the empirical growth data in a hierarchical Bayesian framework (Fig. 1c), where measurement errors are separated from process errors (Carroll et al. 2006; Clark 2007; Muff et al. 2015).

In this study, we address the need to consider alternative species interaction models by comparing the performance of eight models fit to data on the competitive growth of two perennial grass species, *Festuca ovina* and *Agrostis capillaris*, using a Bayesian framework. We start with the Lotka-Volterra competition model and consider three modifications separately and in all combinations: i) addition of frequency dependency, ii) addition of covariance between the early and later growth stages, and iii) modelling of sampling- and measurement errors. The eight different models are compared by their predictive accuracy using the Watanabe-Akaike information criterion.

## Material and Methods

### Competitive growth data

Here, we analyze three years of competitive growth data from ten control plots in a field experiment that was designed to measure the effect of nitrogen addition and an herbicide (glyphosate) on competitive interactions (Daugaard et al. 2011; 2013; 2014). The experiment was established on a former agricultural field with dry, nutrient-poor, sandy soil. The field was fallow for several years prior to the start of the experiment in 2001. The field is quadrangular, surrounded by forest on two sides (south and west) and separated from the neighboring fields by 5-meter broad hedgerows on the other two sides. In 2001, the area was ploughed to 60 cm in order to minimize establishment from the soil seed bank and was prepared for the experiment by harrowing and rolling. Thirty-one grassland plant species covering different life form strategies (CRS strategies *sensu* Grime 2001) were sown in the spring 2001 (Bruus Pedersen et al. 2004). After sowing, plant abundance and species composition were not controlled, except for the removal of woody species (trees and shrubs) every year in the spring. The experiment was set up as a completely randomized block design with ten replicates. Each replicate plot was 7 m x 7 m with a buffer zone of 1.5 m surrounding the plot. A buffer zone of 10 m separated the experiment from the surrounding vegetation. The buffer zones were also sown with the seed mixture. All plots received phosphorus (53 kg/ha), potassium (141 kg/ha), sulphur (50 kg/ha) and copper (0.7 kg/ha) each year.

To study interactions between two perennial grass species, *Festuca ovina* and *Agrostis capillaris*, one permanent 0.5 m x 0.5 m quadrat was placed within each of the plots in June 2007. The quadrat was not placed randomly, but such that both *F. ovina* and *A. capillaris* were clearly abundant. Plant cover and vertical density of all vascular plant species within the quadrats were measured non-destructively by the “pin-point” (also called “point intercept”) method (Kent and Coker 1992) using a pin-point frame with the same dimension as the quadrat and with 25 pin-positions regularly placed at a distance of 10 cm. At each position, a pin with a diameter of 0.5 mm was passed vertically through the vegetation, and the number and species of each contact was recorded. The sampling was performed in the spring and at the end of the growing season for the three-year period 2007-2009.

The pin-point method provides estimates of two important plant ecological variables, plant cover and vertical density, and is well suited for studying the competitive interactions among plant species in natural and semi-natural herbaceous perennial plant communities, where it is difficult to distinguish individual genets (Damgaard 2011; Damgaard et al. 2009). The cover of a specific plant species is defined as the proportion of pins in the grid that touch the species; thus, plant cover measures the cover of the plant species when it is projected onto the two-dimensional ground surface. The vertical density is defined as the number of times a pin touches a specific species, and this measure has been shown to be highly correlated to plant biomass (Jonasson 1983; 1988).

In the subsequent analysis of growth under competition, survival and establishment of *F. ovina* and *A. capillaris*, the data was grouped into three classes: *F. ovina*, *A. capillaris* and an aggregated group of all other vascular plant species.

The data are presented in an electronic supplement (Appendix A).

### Models

The species interactions were analyzed by modelling how the vertical density of *F. ovina*, *A. capillaris* and the other species at the end of the growing season depended on the cover at the beginning of the growing season. The underlying assumption is that the measure of vertical density at the end of the growing season may be used as a measure of population growth as a function of the cover at the beginning of the growing season, growth and competition (Damgaard 2011; Damgaard et al. 2009).

The conceptual competitive growth models for plant communities outlined in eqn. (1) are modified so that they are suited for modelling pin-point cover and vertical density data,

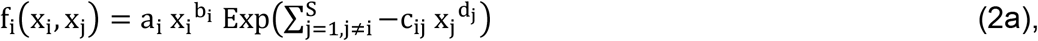

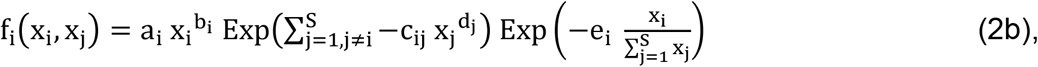

where f_i_(x_i_, x_j_) is the expected vertical density of species i at the end of the growing season, and x_i_ is the cover of species i at the beginning of the growing season. The parameters a_i_ and b_i_ model the expected vertical density as a function of cover in the absence of inter-specific interactions, c_ij_ and d_j_ model the competitive effect of species j on the vertical density of species i (c_ij_ > 0), and e_i_ models the frequency-dependent interaction effect of species i (e_i_ > 0). The parameters b_i_ and d_i_ model possible non-linearity in the effect of cover on vertical density with a domain in the interval (0.5, 2). If e_i_ = 0, there is no evidence of frequency dependence, and model 2b) reverts to the Lotka-Volterra competition model (eqn. 2a).

The measurement errors of cover and vertical density have previously been assumed to be binomial distributed and generalized-Poisson distributed, respectively (Damgaard et al. 2014). However, since we want to model the covariance between the early and later growth stages to account for the possible effect of unmeasured variables (Rinella et al. 2020), both distributions are approximated by standard normal distributions, where (i) the observed pin-point cover measure, y, in a pin-point frame with n pin-positions and an expected cover q is transformed to 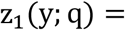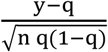, and (ii) the observed pin-point vertical density measure, vd, with an expected value λ and with a species-specific scale parameter ν_i_ is transformed to 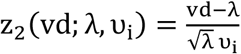.

The measurement error of the standardized cover and vertical density measures is then modelled as

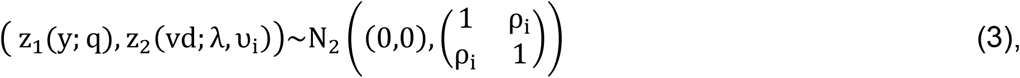

where ρ_i_ is a species-specific correlation coefficient in the domain (−1, 1). If ρ_i_ = 0, then the measurement errors of the standardized cover and vertical density measures are assumed to be uncorrelated, i.e. the covariance between the early and later growth stages is assumed to be accounted for by the competition model and not by additional unmeasured variables.

The effect of measurement errors on the modelling of the inter-specific interactions was investigated by either fixing the latent variables q and λ in eqn. 3 to their empirical mean values (i.e. measurement errors are not modelled), or simulate them during the Bayesian MCMC and thus treating them as parameters along with ν_i_, as is usually done in Bayesian hierarchical modelling (Carroll et al. 2006; Clark 2007; Muff et al. 2015). The structural uncertainty of each species interaction model (eqn. 2a and eqn. 2b) was assumed to be normally distributed and modelled as

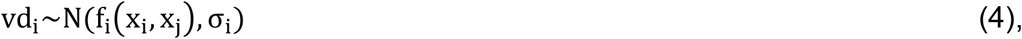

where σ_i_ is the species-specific process error.

In total, 2^3^ = 8 different models were fitted to the data by combing a Lotka-Volterra competition model with or without (1) frequency dependency, (2) including the effect of confounding factors, and (3) including the effect of measurement errors in all possible combinations.

### Estimation

We have chosen to treat the three years of competitive growth data from each of the ten control plots as independent events because i) the within-plots among-year covariation is expected to be minimal since we model plant growth of each year, and ii) considering plot as a random effect is not compatible with modelling covariance between early and late plant growth.

The joint Bayesian posterior probability distribution of the parameters in the eight models was calculated using Bayesian Markov Chain Monte Carlo (MCMC; Metropolis-Hastings) simulations of 100,000 iterations with a burn-in period of 70,000 iterations and normal candidate distributions (Carlin and Louis 1996). The prior probability distributions of all parameters and latent variables were assumed to be uniformly distributed in their specified domains under the additional constraints that c_ij_ < 5, e_i_ < 5, 0.5 < ν_i_ < 5, except for *σ*_i_ which was assumed to be inverse gamma distributed with parameters (0.1, 0.1).

Plots of the deviance and trace plots of all parameters were inspected in order to check the fitting and mixing properties of the used sampling procedure. Residual plots of the vertical density for each species were inspected to check the fitting properties of the different models. The statistical inferences were assessed using the calculated credible intervals, i.e. the 95% percentiles of the marginal posterior distribution of the parameters.

The eight different models were compared by their predictive accuracy using the Watanabe-Akaike information criterion (WAIC, Gelman et al. 2014; McElreath 2016; Watanabe 2010). Using WAIC, the effective number of parameters, a measure of how many hypothetical independent (uncorrelated and unbounded) parameters the model uses, was also estimated for each model.

All calculations were done using Mathematica (Wolfram 2019). The software code and all results are given as an electronic supplement (Appendix B).

## Results

The MCMC iterations of all eight models converged with acceptable mixing properties (Appendix B). Based on the residual plots (Fig. S1), the fit of all models was judged to be acceptable. A summary of the marginal posterior probability distributions of all parameters are shown in Table S1. The predictive accuracy of the eight models using WAIC (Table 1) was very similar to that calculated by the closely-related Deviance Information Criterion (Spiegelhalter et al. 2002; Appendix B). Agreement between predictive accuracy and relatively low structural uncertainty of the models (Table S1, σ_i_) was satisfactory.

**Table 1.** Predictive accuracy and the estimated number of effective parameters using the Watanabe-Akaike information criterion (WAIC). 2^3^ = 8 different models were fitted to the data by combining a Lotka-Volterra competition model without frequency dependency (eqn. 2a) or with frequency dependency (eqn. 2b), including the effect of unmeasured variables, and including the effect of measurement errors in all possible combinations. The best predictive models are the ones with the lowest WAIC value.

| Model number | Frequency dependence | Unmeasured variables | Measurement errors | WAIC | Effective number of parameters |
| --- | --- | --- | --- | --- | --- |
| 1 | – | – | – | 935.89 | 13.86 |
| 2 | + | – | – | 925.99 | 13.25 |
| 3 | – | + | – | 851.05 | 11.23 |
| 4 | + | + | – | 849.88 | 14.00 |
| 5 | – | – | + | 705.20 | 83.39 |
| 6 | + | – | + | 594.80 | 92.01 |
| 7 | – | + | + | 692.06 | 83.84 |
| 8 | + | + | + | 700.73 | 90.44 |

Based on predictive accuracy, the model that best fit the competitive growth data was one in which frequency dependency was added to the Lotka-Volterra competition model (eqn. 2b) and where the measurement error was included, but not a potential effect of unmeasured variables (Table 1, WAIC = 594,80). This was also the model with lowest structural uncertainty (Table S1, σ_i_).

The four models with the highest predictive accuracy were those that included the effect of measurement errors, even though they also had the highest number of effective parameters (Table 1, the four models with lowest WAIC). However, if measurement errors were ignored, then the model with the highest predictive accuracy was again one in which frequency-dependence was added to the Lotka-Volterra competition model, but then the predictive accuracy was increased when the effect of unmeasured variables was included (Table 1, WAIC = 849.88). Consequently, the result that the predictive accuracy was increased when frequency-dependency was added to the Lotka-Volterra competition seemed robust in this case of competitive growth between two grass species.

As explained above, the effect of not taking unmeasured variables into account is that the competitive effect may be biased, but this was only significant in one out of 24 cases (Table S1; *c*_23_ figures in bold), which is in agreement with random expectations. However, the credibility intervals of the competition coefficients were relatively wide, so the statistical power is correspondingly low.

The predicted effect of interspecific competition differed significantly among the eight different models even though they were fitted to the same data (Fig. 2). The mean expected vertical density of *F. ovina* as a function of the cover of itself and *A. capillaris* when fitted with the eight different models was consistently lower in the four models where the measurement error was included in a hierarchical model (Fig. 2), and these were also the models that received most support from the data (Table 1).

**Fig 2.**
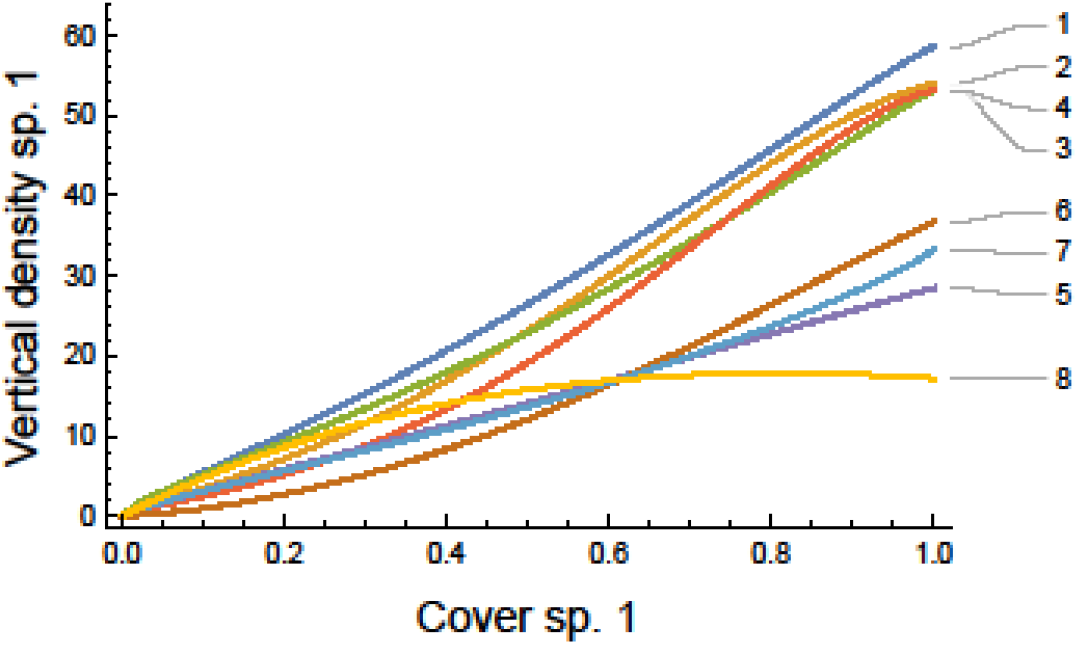
The mean expected vertical density (per frame) of species 1 (*Festuca ovina*) as a function of the cover of species 1 and species 2 (*Agrostis capillaris*) when fitted with the eight different models (the model number corresponds to the model # in Table 1). The combined cover of species 1 and species 2 is set to one; eqn. 2: *f*_1_(*x*_1_, *x*_2_, *x*_3_) = *f*_1_(*x*_1_, 1 − *x*_1_, 0).

## Discussion

This case study provides a compelling example of the need to consider somewhat more complex competition models than the standard Lotka-Volterra model in empirical studies of species interactions in plant communities. While the use of hierarchical models to account for measurement errors is becoming the new standard for statistical modelling, most empirical species interaction studies are still modelled without considering frequency-dependency or the effects of unmeasured variables.

The use of the classical Lotka-Volterra model for species interactions has been criticized by several authors. Some researchers have argued that the neutral model (Hubbell 2001) is a more relevant null-model for species interaction, but neutrality or the absence of species interactions can be considered a degenerate case of the Lotka-Volterra model when *α*_*ij*_ (eqn. 1a) is zero, and this null-hypothesis is in practice tested indirectly. Interestingly, Neill et al. (2009) constructed a “null-model” that resembles the dynamics of frequency-dependent species interaction models using only maximum entropy assumptions. The approach chosen here is to assume that the simplest hypothesis, or null-hypothesis (H_0_), is neutrality, which is tested indirectly using the Lotka-Volterra model. The next hypothesis (H_1_) is the linear effect of density on plant growth, which is modelled by Lotka-Volterra type models. This hypothesis is followed by more complicated species interactions models with higher order terms (H_2_), in which frequency-dependent species interaction models are a special case. There are a number of possible species interaction models with higher order terms or other modifications of the standard species interaction approach, e.g. (i) if the outcome of competitive interactions depends on the local spatial distribution, i.e. whether the species are randomly distributed or aggregated (Bolker and Pacala 1999; Damgaard 2004; Stoll and Prati 2001), or (ii) when more species interact, e.g. in a rock -paper -scissors type of interaction (Levine et al. 2017) or (iii) when intraspecific genetic variation may play a role in the outcome of species interactions (Ehlers et al. 2016).

In the present case study of interspecific interactions between two grass species, we found that species interaction models that included frequency-dependency were better supported by the data and improved the prediction accuracy. This finding is in agreement with the study of Harpole and Suding (2007), who found relatively strong frequency-dependency among four annual plant species. These findings are relevant to the recent emphasis on species-specific soil-plant interactions as a potentially important and general mechanism for regulating plant populations (Heinen et al. 2020; Mazzoleni et al. 2015a; Mazzoleni et al. 2015b; van der Putten et al. 2013).

There was only limited correlation between early and later growth stages, which may indicate that the observed growth was not influenced by unmeasured variables that affected both early and later growth stages or because the experiment was replicated (Rinella et al. 2020). In other empirical competition studies, unmeasured variables have been shown to influence the estimated competition coefficients (Rinella et al. 2020).

Although the effective number of parameters was relatively high when measurement errors were included, the four models with the highest predictive accuracy were those that included the effect of measurement errors. These additional effective parameters were probably mainly used in describing measurement uncertainty and not in the modelling of the competitive interactions. It has been demonstrated that if the effects of measurement errors are ignored, model predictions may be biased (Carroll et al. 2006; Damgaard 2020; Detto et al. 2019) so it is therefore advisable to consider the possible effect of measurement errors when modelling species interactions.

If the aim of an empirical study is to make predictions of plant community dynamics, it is possible to increase the overall predictive performance of models by using the calculated WAIC for model averaging (McElreath 2016).

In the present study, there were only two-yearly sampling times, but if there had been more, it would have been interesting to analyze whether the growth stage of the plant species had an effect on which species interaction model was best supported by the growth data.

Theoretical studies of plant community dynamics and the question of how many plant species may coexist in plant communities have been dominated by Lotka-Volterra type competition models (Barabás et al. 2018; Chesson 2000; Damgaard 2005). These models are characterized by a relatively low number of species at ecological equilibrium, and much theoretical research has been motivated by the need to explain the apparent paradox of the relatively many observed plant species compared to the number that is expected according to Lotka-Volterra type competition models (Hutchinson 1961). If frequency-dependent mechanisms play an important role in the regulation of plant populations, then the expected number of species at ecological equilibrium will be higher and may explain the observed relatively high frequency of rare species (Enquist et al. 2019). Consequently, frequency-dependence needs to be integrated into theoretical, as well as empirical, plant competition models.

Spatial dynamics have also been used to explain the high number of apparently coexisting plant species. Theoretical studies (e.g. May and Nowak 1994; Tilman 1994) have suggested that if there is a trade-off between density-independent fitness components (ex. viability, fecundity, colonizing ability) and competitiveness, then an infinite number of species could coexist at equilibrium. However, Adler and Mosquera (2000) have challenged this conclusion by generalizing the competitiveness function. They showed that with a biologically realistic, smooth competitiveness function, only a few species would be able to coexist at equilibrium despite such a trade-off.

To paraphrase Einstein “The model should be as simple as possible and as complex as necessary”. Alternative models evaluated with Bayesian statistical methods allow us to apply Einstein’s dictum to plant community dynamics. Our results suggest that in ecology, more complexity than we might wish for is necessary if we are to understand and predict competitive processes in plant communities.

## Supporting information

Electronic supplements

## Acknowledgements

The data were collected in a project supported by Miljøstyrelsen (the Danish EPA).

## Electronic Supplements

Appendix A: Pin-point data (Excel)

Appendix B: Software code (Mathematica notebook)

Table S1: Summary of marginal posterior distributions of parameters (Excel file)

Fig. S1: Residual plots of vertical density

