## Supplementary material for "The need for alternative plant species interaction models": Electronic supplements: Figure S1.docx

Figure S1. Residual plots. Histogram and Quantile plots for the compactness of the three species

*Festuca ovina	Agrostis capillaris*	Other species

Model 1

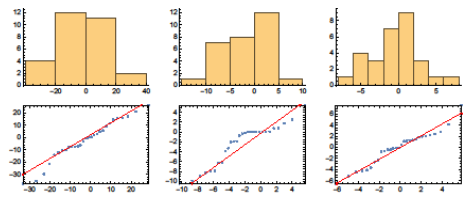

Model 2

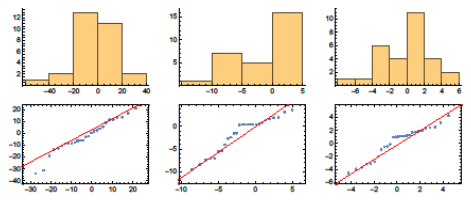

Model 3

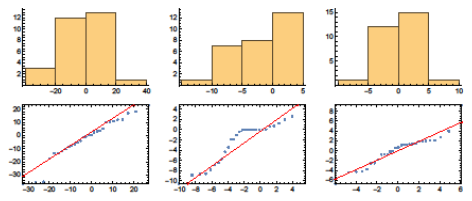

Model 4

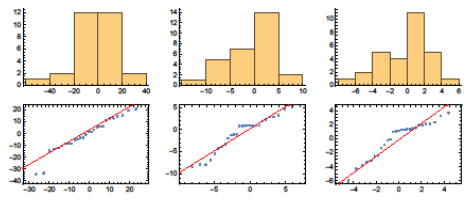

*Festuca ovina	Agrostis capillaris*	Other species

Model 5

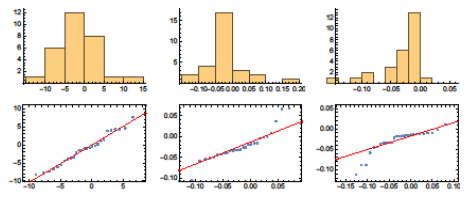

Model 6

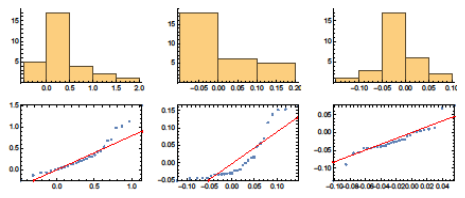

Model 7

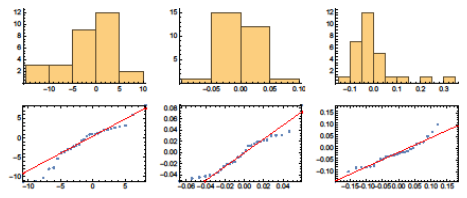

Model 8

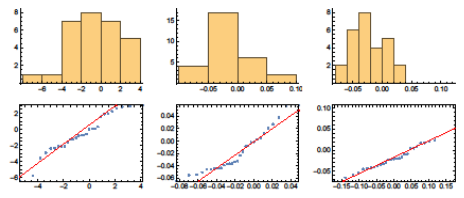
